# A Genome Scan for Genes Underlying Adult Body Size Differences between Central African Pygmies and their Non-Pygmy Neighbors

**DOI:** 10.1101/187369

**Authors:** Trevor J. Pemberton, Paul Verdu, Noémie S. Becker, Cristen J. Willer, Barry S. Hewlett, Sylvie Le Bomin, Alain Froment, Noah A. Rosenberg, Evelyne Heyer

## Abstract

**Background:** Central African hunter-gatherer Pygmy populations have reduced body size compared with their often much larger agricultural non-Pygmy neighbors, potentially reflecting adaptation to the anatomical and physiological constraints of their lifestyle in tropical rainforests. Earlier studies investigating the genetics of the pygmy phenotype have focused on standing height, one aspect of this complex phenotype that is itself a composite of skeletal components with different growth patterns. Here, we extend the investigations of standing height to the variability and genetic architecture of sitting height and subischial leg length as well as body mass index (BMI) in a sample of 406 unrelated West Central African Pygmies and non-Pygmies.

**Results:** In addition to their significantly reduced standing height compared with non-Pygmies, we find Pygmies to have significantly shorter sitting heights and subischial leg lengths as well as higher sitting/standing height ratios than non-Pygmies. However, while male Pygmies had significantly lower BMI compared with male non-Pygmies, the BMI of females were instead similar. Consistent with prior observations with standing height, sitting height and subischial leg length were strongly correlated with inferred levels of non-Pygmy genetic admixture while BMI was instead weakly correlated, likely reflecting the greater contribution of non-genetic factors to the determination of body weight compared with height. Using 196,725 SNPs on the Illumina Cardio-MetaboChip with genotypes on 358 Pygmy and 169 non-Pygmy individuals together with single-and multi-marker association approaches, we identified a single genomic region and seven genes associated with Pygmy/non-Pygmy categorization as well as 9, 10, 9, and 10 genes associated with standing and sitting height, sitting/standing height ratio, and subischial leg length, respectively. Many of the genes identified have putative functions consistent with a role in determining their associated trait as well as the complex Central African pygmy phenotype.

**Conclusions:** These findings highlight the potential of modestly sized datasets of Pygmies and non-Pygmies to detect biologically meaningful associations with traits contributing to the Central African pygmy phenotype. Moreover, they provide new insights into the phenotypic and genetic bases of the complex pygmy phenotype and offer new opportunities to facilitate our understanding of its complex evolutionary origins.

## Background

Central African hunter-gatherer populations have historically been called “Pygmies” in reference to their proportionally reduced body size compared with their agricultural non-Pygmy neighbors [1]. Among Pygmy males, mean adult standing height ranges from 142 cm among the Efe from eastern Democratic Republic of Congo (DRC; 142 cm) to 161 cm among the Twa from western DRC (161 cm), averaging 150.6 cm across 23 Central African Pygmy populations (standard deviation [SD]=6.7 cm) [2,3]. This average is notably shorter than the average adult standing height of 167.3 cm among males from 252 Sub-Saharan non-Pygmy populations (SD=5.7 cm) [3]. Nevertheless, no discontinuity in adult standing height exists among Pygmies and non-Pygmies, and contemporary populations are categorized as Pygmy based on both cultural criteria and adult standing height [1,3-7].

It has been hypothesized that the diminutive body size of Central African Pygmies is the outcome of adaptive processes in response to hunter–gatherer lifestyles in tropical rainforests with high levels of pathogen exposure [8]. Hypotheses for the basis of adaptation have considered morphological adaptation to thermoregulation in a hot and humid environment [9], metabolic costs of hunting and gathering in an environment in which food is scarce [10-12], and an early onset of reproduction in a context of high mortality rates [13]. Physiological studies in both West-and East-Central African Pygmies have found altered glucose homeostasis, insulin secretion, and free fatty-acid profiles in the presence of normal human growth hormone (HGH) levels [14,15]. In addition, reduced growth rates during the first two years of life [16] as well as adolescence [17,18] compared with non-Pygmies have been reported, possibly reflecting perturbation of the insulin-like growth factor 1 (IGF1) [19,20] and HGH [21] receptor signaling pathways [22]. Limitations on physiological data, however, together with a paucity of demographic, epidemiological, and paleoanthropological data for Central Africa have contributed to the importance of genetics for assessing hypotheses about contemporary pygmy body size and its evolution [23].

Population-genetic studies have inferred a probable common ancestral origin for West Central African Pygmy populations ~3,000 years ago and for West-and East-Central African Pygmy populations ~25,000 years ago, following their divergence from ancestral Central African non-Pygmy populations ~50,000-70,000 years ago [24-28]. Substantial levels of non-Pygmy genetic admixture have been observed across Central African Pygmy populations [24-27,29-31], correlating positively with adult standing height [32-34]. The general genetic difference between Pygmies and non-Pygmies together with the correlation of genetic admixture and standing height suggests that adult body size differences among Central African Pygmies and neighboring non-Pygmies are attributable in large part to genetic factors, arguing against a view that diminutive Central African pygmy body size is the consequence solely of phenotypic plasticity in a challenging nutritional and parasitic environment [8].

Over the past ten years, numerous genome-wide association studies (GWAS) have sought to identify genetic factors determining adult standing height [35-51], a high-heritability trait (≥69% [52-54]) despite being strongly influenced by environmental factors [55,56]. The largest study to date identified 697 common single-nucleotide polymorphisms (SNPs) significantly associated with adult standing height in a cohort of ~250,000 individuals of recent European ancestry that together explained 16% of the variance and 20% of the heritability of adult standing height in their cohort [51]. Recently, two GWAS on separate admixed Afro-American cohorts identified novel loci significantly associated with adult standing height [57,58], illustrating the potential of populations with a component of recent African ancestry to contribute new information to the study of adult standing height. In this context, Central African Pygmy and non-Pygmy populations—with marked differences in adult standing height that are strongly correlated with levels of genetic admixture—provide a potentially powerful framework within which to understand the genetic basis of adult human height.

A number of recent studies explored the genetic basis of adult standing height variation in Central Africans, comparing genome-wide SNP data on ≤3 Pygmy populations and their non-Pygmy neighbors [33,34]. These studies uncovered signatures of differential polygenic adaptation between Pygmies and non-Pygmies primarily in genomic regions containing genes associated with immunity and metabolism, as well as evidence of differential signatures in Western and Eastern Pygmies suggesting a partially convergent origin of the pygmy phenotype in these groups. Perry *et al*. [34] further identified genetic associations with standing height that included four regions previously implicated in adult standing height variation in Europeans. However, no association was detected with the ~100 European standing-height-associated SNPs present in these modestly sized datasets of 67–169 Pygmies and 58–61 non-Pygmies, strongly suggesting that the differing stature of Pygmies and non-Pygmies is attributable at least in part to genetic factors that have arisen since the ancestral split between Central Africans and Europeans.

While these recent studies have focused on the differing standing height of Central African Pygmies and non-Pygmies, other morphological differences also exist between these two groups. For example, Pygmy body weight appears proportionally more reduced relative to standing height compared with non-Pygmies [59,60], a difference that cannot be wholly explained by differential nutrition levels in Pygmies and non-Pygmies [61]. In the context of the differential diets of hunter-gatherer Pygmies and agriculturalist non-Pygmies, and the greater seasonal variation in food availability experienced by hunter-gatherers compared with agriculturalists, such a difference might reflect the coevolution of differences in energy usage and storage in response to their different lifestyles. In addition, craniofacial and skeletal dissimilarities are also evident, including skull morphology [59,62] as well as leg and forearm length [59,63] where Pygmies generally have shorter legs and forearms relative to trunk length compared with non-Pygmies, potentially reflecting evolutionary changes in shared endoskeletal development pathways in response to the anatomical constraints of Pygmy hunting and gathering activities in the tropical rainforest. The reported dissimilarities in trunk and leg length patterns accord with their differential growth patterns in response to nutritional [64,65] and health [64] factors that vary between hunter-gatherer Pygmies and agriculturalist non-Pygmies [66-68], and are consistent with the view that both common and distinct pathways contribute to the determination of upper and lower body size [44]. In this context, while genes contributing to both trunk and leg length determination may be detected by genetic studies investigating standing height variation in Central Africans, those distinct to each trait may not because of the confounding effects of their variable contributions to standing height across groups. Thus, a joint analysis of variability in trunk and leg lengths in Pygmies and non-Pygmies might disentangle their relative contributions to Pygmy short stature and provide a more complete picture of the genetic architecture and anatomical constraints underlying the diminutive body size of contemporary Central African Pygmies.

Here, using body size and weight measurements available for individuals from seven Pygmy and three non-Pygmy populations from West Central Africa (**Figure 1**; **Table 1**) together with a genome-wide SNP data, we investigate the genetic basis of standing and sitting height, and subischial leg length in Central Africans. Our sample set includes 358 Pygmy and 169 non-Pygmy individuals genotyped on the Illumina Cardio-MetaboChip [69], a SNP genotyping microarray that interrogates a set of 68,126 SNPs previously identified in well-powered GWAS conducted by the Body Fat Percentage [70], CARDIoGRAM (coronary artery disease and myocardial infarction) [71] DIAGRAM (type 2 diabetes) [72], GIANT (anthropometric traits) [45,73,74], Global Lipids Genetics (lipids) [75], HaemGen (hematological measures) [76], ICBP (blood pressure) [77], MAGIC (glucose and insulin) [78-80], and QTIGC (QT interval) [81,82] consortia in addition to 146,453 SNPs chosen to facilitate fine-mapping of the genomic regions surrounding these 68,126 SNPs. The Cardio-MetaboChip therefore allows us to both investigate potential associations between the 1,050 SNPs associated with adult standing height variation [45] in Europeans with adult standing height variation in Central Africans. Moreover, it will allow us to explore Pygmy/non-Pygmy genetic differences in genes previously implicated in the determination of anthropometric, metabolic, and physiometric traits that we would expect to have been subject to historical natural selection under the hypotheses proposed for the evolution of the diminutive body size of Central African Pygmies [8-13].

**Table 1.** Populations and their sample sizes.

| Category | Population |  | Country | Trait data | Sample size |  |
| --- | --- | --- | --- | --- | --- | --- |
|  | Code | Name |  |  | Total | Genetic dataset |
| Pygmies | AKA | Aka | CAR | No | 26 | 21 (M=14; F=7) |
|  | AKM | Aka (Mbatì) | CAR | No | 17 | 8 (M=6; F=2) |
|  | CBK | Central/Eastern Baka | Cameroon | Yes | 36 | 29 (M=14; F=15) |
|  | EBK | Southeastern Baka | Cameroon | Yes | 11 | 10 (M=10; F=0) |
|  | SBK | Southern Baka | Cameroon | No | 50 | 41 (M=16; F=25) |
|  | BZN<sup>a</sup> | Northern Bezan | Cameroon | Yes | 27 | 27 (M=12; F=15) |
|  | BZS<sup>a</sup> | Southern Bezan | Cameroon | Yes | 48 |  |
|  | CBG | Central Bongo | Gabon | Yes | 24 | 21 (M=14; F=7) |
|  | EBG | Eastern Bongo | Gabon | Yes | 30 | 19 (M=13; F=6) |
|  | SBG | Southern Bongo | Gabon | Yes | 34 | 27 (M=13; F=14) |
|  | KOY | Koya | Gabon | Yes | 25 | 19 (M=12; F=7) |
|  | KOL | Kola | Cameroon | No | 31 | 24 (M=18; F=6) |
|  | NSU | Nsua (Efe) | Uganda | No | 17 | 11 (M=6; F=5) |
| **Total** |  |  |  |  | **376** | **257 (M=148; F=109)** |
| Non-Pygmies | AKL | Akele | Gabon | No | 12 | 8 (M=7; F=1) |
|  | CFG | Fang | Cameroon | No | 36 | 33 (M=19; F=14) |
|  | NZI | Nzime | Cameroon | Yes | 23 | 20 (M=3; F=17) |
|  | NGB | Ngumba | Cameroon | No | 58 | 43 (M=20; F=23) |
|  | TIK | Tikar | Cameroon | Yes | 20 | 20 (M=12; F=8) |
|  | BGD | Bangando | Cameroon | Yes | 6 | 6 (M=3; F=3) |
|  | BKJ | Konjo | Uganda | No | 27 | 19 (M=8; F=11) |
| **Total** |  |  |  |  | **182** | **149 (M=72; F=77)** |

Numbers of males (M) and females (F) appear in parentheses.
<sup>a</sup>The two Bezan populations were merged into a combined BEZ population in the 406UNRELAT dataset.

**Figure 1.**
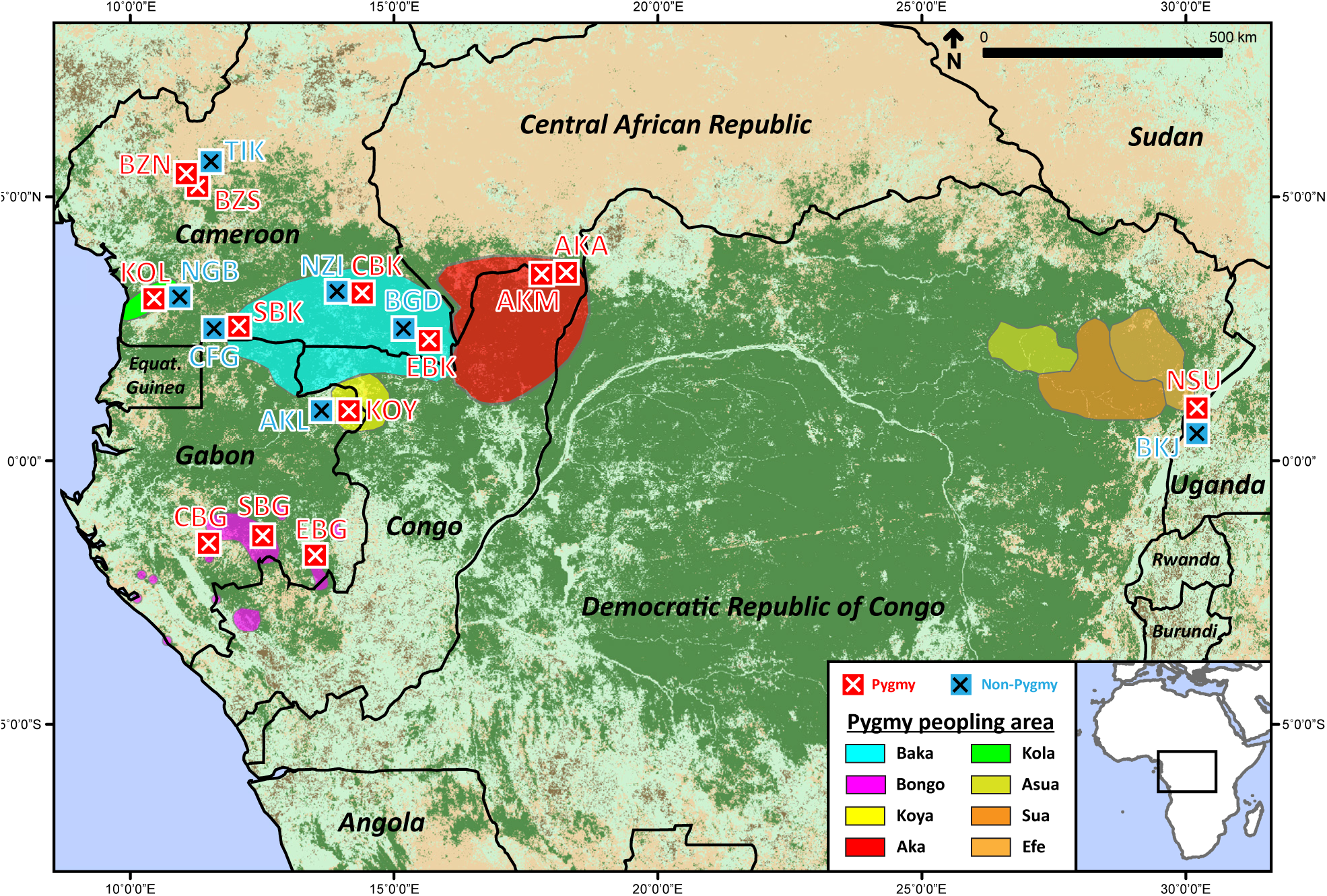
Sampling locations. Pygmy/non-Pygmy categorization was assessed in ethnographic field work, relying on numerous cultural criteria that do not include adult height (see **Material and Methods**). Population codes follow **Table 1**. Population ranges were inferred from ethnographic field work [7].

Using a mixed-effect variance components model to correct for population stratification together with a multi-marker gene-based association method, we investigate the genetic determination of adult standing and sitting height, subischial leg length, and the ratio of sitting to standing height to identify genes and variants specific to their variation in Pygmies and in non-Pygmies. We identify three genomic regions containing SNPs with significant allele-frequency differences between Pygmies and non-Pygmies, and numerous genes significantly associated with each considered trait of which many have known functions compatible with a deterministic role in Central African body size. Our results shed new light on the complex genetic architecture of the Central African pygmy phenotype, and highlight novel pathways contributing to its height and metabolic aspects that provide new opportunities for more comprehensive comparisons of Pygmies and non-Pygmies.

## Results

Our objective was to perform a GWAS to advance our understating of the genetic basis of phenotypic differences between Central African Pygmies and their non-Pygmy neighbors, categorized as such based solely on cultural criteria and *not* on adult standing height (see **Materials and Methods**). Unlike normal GWAS where homogeneity among samples is desirable, we were instead conducting a GWAS in a more complex situation where significant phenotypic and genetic differences that provide potentially greater power to detect biologically meaningful associations are to be expected. We therefore first had to establish the levels phenotypic and genetic diversity that existed in our sample set to facilitate the design of the association analyses that we will perform.

### Differences in measured traits between Pygmies and non-Pygmies

We first investigated whether the height and weight traits with measurements available for our sample set exhibited significant differences between Pygmies and non-Pygmies. Of the 527 individual in our sample set, we had trait measurements available for 115 Pygmy and 44 non-Pygmy individuals representing seven Pygmy and three non-Pygmy populations from West Central Africa (Additional File 1: **Table S1**).

Among the 115 Pygmies in our sample set, average male standing height was 155.67 cm (SD=6.55) while average female standing height was 148.95 cm (SD=5.82), in contrast to the 44 non-Pygmies whose average male and female standing heights were 167.21 cm (SD=4.61) and 154.45 cm (SD=6.23), respectively (**Figure 2A**). These values are consistent with average standing heights reported by prior studies of Central African Pygmy (males=154.85 cm with SD=3.08 and females=146.30 cm with SD=2.87) [3,9,83-88] and non-Pygmy (males=164.98 cm with SD=2.36 and females=155.96 cm with SD=0.85) [9,86,87,89] populations [59] and highlight the significant difference in stature that exists between Pygmies and non-Pygmies (*P*_males_=6.13×10^−8^ and *P*_females_=2.65×10^−4^; Wilcoxon rank-sum test).

**Figure 2.**
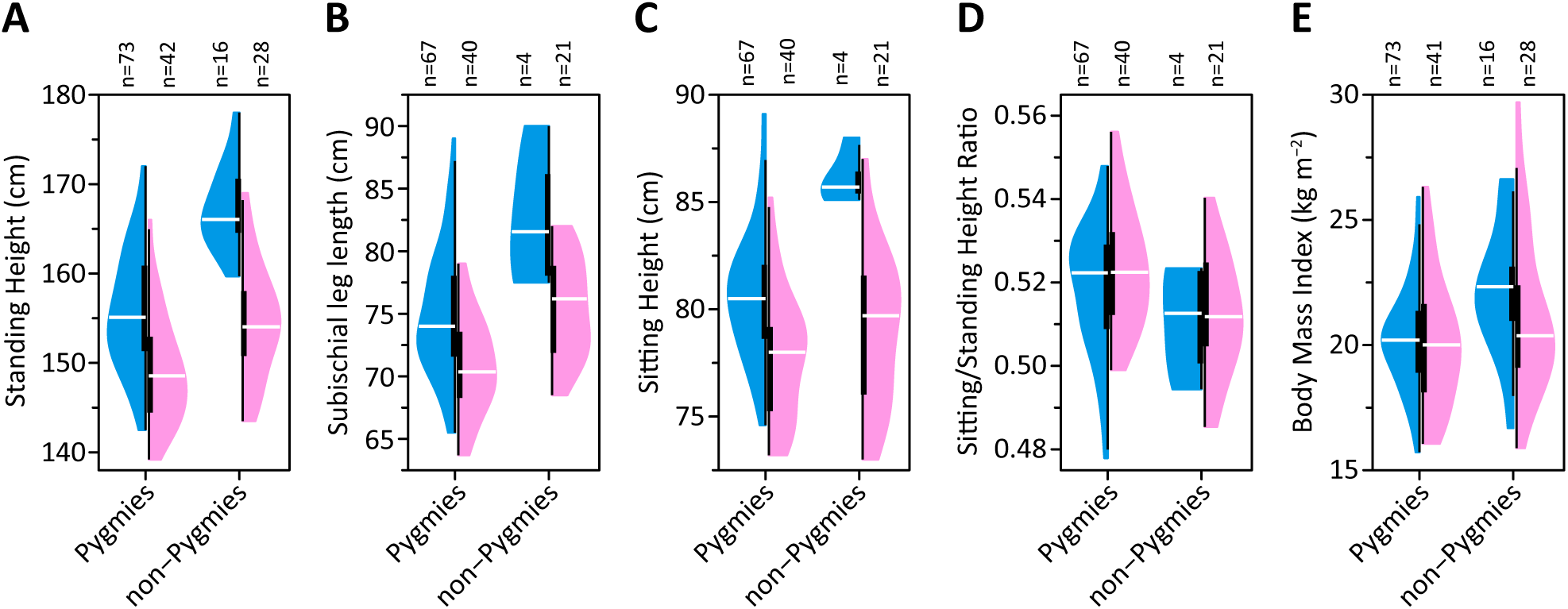
Distributions of standing height, sitting/standing height ratios, and BMI in Pygmies and non-Pygmies. Split violin plot [146] representations of the distributions of (**A**) standing heights, (**B**) subischial leg lengths, (**C**) sitting heights, (**D**) sitting/standing height ratios, and (**E**) BMI shown separately for males (blue) and females (pink). The number (*n*) of males and females with data available for each measurement is provided above each plot. The mean and SD of each group are reported in **Table S1** (Additional File 1). Each “violin” contains a vertical black line (25%–75% range) and a horizontal white line (median), with the width depicting a 90°-rotated kernel density trace.

Consistent with the high correlations expected between standing height and other height-related traits (Additional File 2: **Figure S1**), subischial leg lengths of male (mean=74.97 cm, SD=4.91) and female (mean=70.81 cm, SD=3.84) Pygmies were significantly shorter than those of male (mean=82.65 cm, SD=5.85) and female (mean=75.44 cm, SD=3.97) non-Pygmies (*P*_males_=0.007 and *P*_females_=9.38×10^−5^; **Figure 2B**). However, while the sitting heights of male Pygmies (mean=80.62 cm, SD=2.92) were significantly shorter than those of male non-Pygmies (mean=86.13 cm, SD=1.28; *P*=0.002), sitting heights of female Pygmies (mean=77.74 cm, SD=2.89) were instead similar to female non-Pygmies (mean=78.96 cm, SD=3.51; *P*=0.073; **Figure 2C**). These findings are consistent with leg length contributing more to the differential stature of Pygmies and non-Pygmies than upper body (trunk and head) length [59]. If we instead compare the ratio of sitting to standing height between Pygmies and non-Pygmies (**Figure 2D**), we observe significantly higher ratios among Pygmies than among non-Pygmies for females (*P*=0.003) but not for males (*P*=0.203) despite male and female Pygmies exhibiting a similar shift toward higher ratios compared with non-Pygmies (**Figure 2D**). The lack of significance for males likely reflects our decreased power to detect such a difference due to only four male non-Pygmies having both sitting and standing height data available compared with 67 male Pygmies.

Focusing on BMI instead of body weight since it accounts for the differential stature of Pygmies and non-Pygmies (**Figure 2A**), values were similar among male (mean=20.16, SD=1.95) and female (mean=20.06, SD=2.56) Pygmies (*P*=0.587), while those of male non-Pygmies (mean=22.31, SD=2.63) were significantly higher than those of female non-Pygmies (mean=20.16, SD=3.07; *P*=0.031; **Figure 2E**). Consequently, Pygmies had significantly lower BMI than non-Pygmies when considering males (*P*=2.74×10^−4^) but not females (*P*=0.186). While our BMI are consistent with those reported by prior studies for male and female Pygmies in 26 populations (mean=20.04 with SD=0.80 and mean=20.33 with SD=0.90, respectively) and male non-Pygmies in 10 populations (mean=21.48, SD=0.90), our female non-Pygmy BMI are notably lower than those of earlier studies (mean=21.65, SD=1.02) [59].

These findings reaffirm earlier observations of significant differences in standing height between Central African Pygmies and non-Pygmies, and highlight apparent differences in the contribution of trunk and leg length to these differences. However, our findings with BMI contrast with patterns in prior studies that indicate differences exist between Pygmies and non-Pygmies for both males *(P*=1.21×10^−4^, Wilcoxon rank sum test of population means) and females (*P*=4.57×10^−4^) and no male-female differences among Pygmies (*P*=0.858, Wilcoxon signed rank test) and non-Pygmies (*P*=1).

### Genetic structure patterns

Now that we had established that significant differences existed between Pygmies and non-Pygmies for the height-related measurements available in our sample set, we next investigated the levels of genetic differentiation that are present among our Pygmy and non-Pygmy samples. Our genetic dataset consisted of 406 unrelated individuals from 20 populations with genotypes at 153,798 SNPs included on the Illumina Cardio-MetaboChip following the removal of lower quality SNPs and related individuals (see **Material and Methods**). In agreement with earlier studies [24-27,29,30,33], multidimensional scaling (MDS) analysis of pairwise allele-sharing dissimilarities (ASD) among these 406 unrelated individuals supported three main components to genetic differentiation among Central Africans (**Figure 3A**). First, differentiation between Pygmies and non-Pygmies— categorized as such based on cultural criteria and not on height—largely determined the first MDS dimension. Second, though non-Pygmies formed a single tight cluster in the first two MDS dimensions reflecting the recent history of these mostly Bantu-speaking populations [90-94], differentiation among Pygmy populations was evident in the second dimension, driven partly by differences between the West African Baka (CBK, EBK, SBK) and Aka (AKA, AKM) populations and the East African Nsua (NSU). Dispersion among Pygmies—measured as the variance among their ASD values—was significantly higher than among non-Pygmies (2.07×10^-5^ and 7.76×10-^6^, respectively; *P*<10^-16^, one-sided *F*-test), consistent with appreciable reproductive isolation following their ancestral divergences over the last ~25,000 years [24-26,92,95,96]. Third, the degree of genetic differentiation from non-Pygmies varied among Pygmy populations, consistent with what might be expected in the presence of variable levels of non-Pygmy gene flow into Pygmy populations. The Bezan, Kola and Bongo Pygmies clustered closer to the non-Pygmies (mean ASD=0.2047, SD=0.0027, across 20,413 pairs of individuals) than did the Baka and Aka (mean ASD=0.2051, SD=0.0025, 16,241 pairs) and Nsua (mean ASD=0.2072, SD=0.0036, 1,639 pairs) Pygmies; the first and third quantiles of the full distribution of ASD values were 0.1983 and 0.2053, respectively.

**Figure 3.**
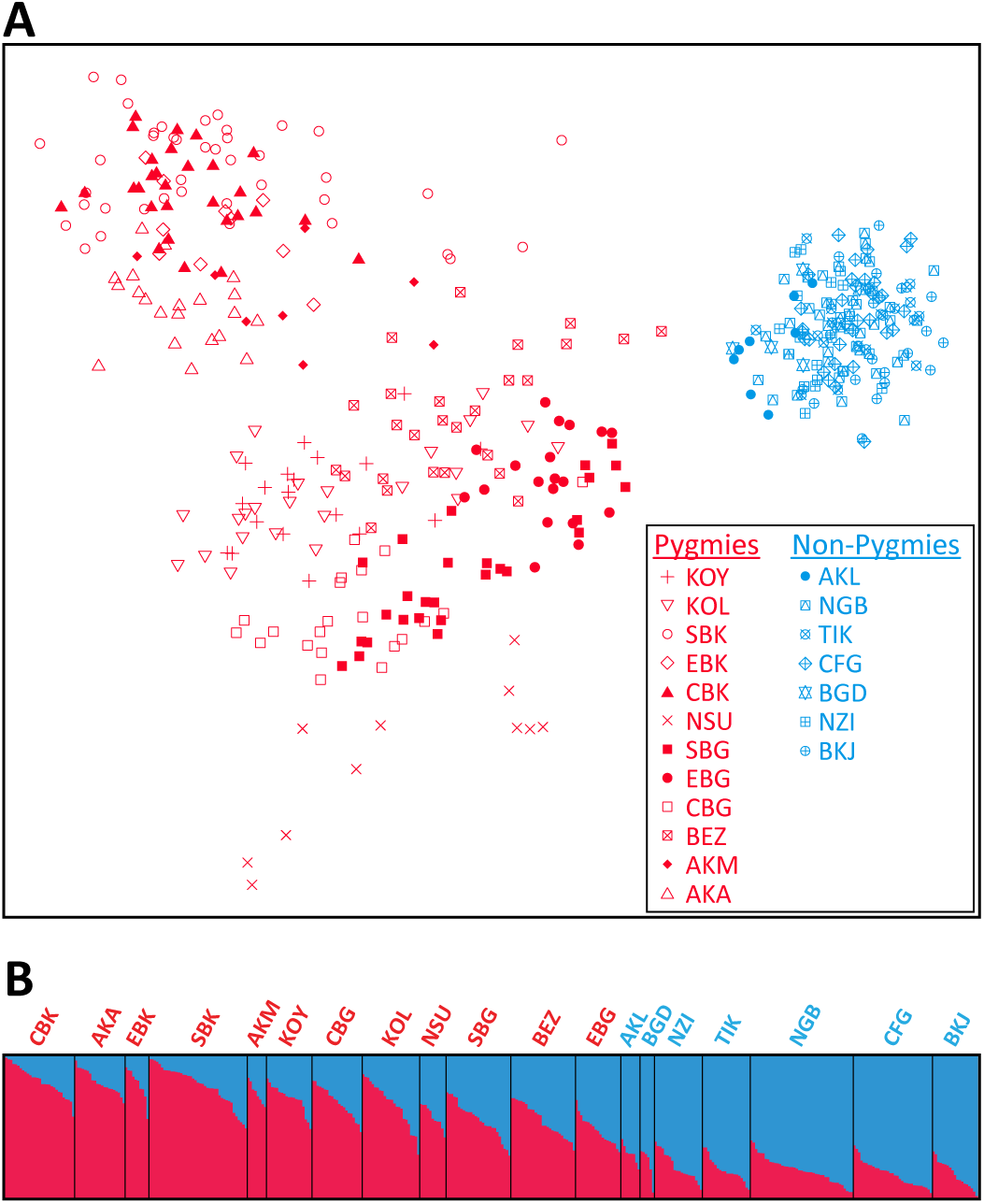
Genetic diversity and admixture patterns. **(A)** A two-dimensional multidimensional scaling (MDS) representation of allele-sharing distances (ASD) for 406 unrelated individuals in the genetic dataset. The first dimension is plotted on the horizontal axis, and the second dimension on the vertical axis. The Spearman *ρ* between pairwise Euclidean distances in the MDS plot and ASD matrix is 0.478. **(B)** Population structure inferred by *STRUCTURE* at *K*=2 for 406 unrelated individuals. Each individual is shown as a thin vertical line partitioned into two components representing inferred genotype membership proportions in the two clusters. Black vertical lines separate individuals from different populations. The most salient population structure observation is the Pygmy/non-Pygmy distinction at *K*=2.

### Genetic admixture patterns

To further explore the presence of potential signals of variable genetic admixture between Central African Pygmies and non-Pygmies suggested by our MDS analysis, we applied the Bayesian clustering algorithm implemented in *STRUCTURE* [97,98] to four non-overlapping sets of 10,106 low-linkage-disequilibrium (LD) genome-wide SNPs. At *K*=2 (**Figure 3B**), Pygmies were assigned greater membership in one cluster (“red”) while non-Pygmies were assigned greater membership in the other cluster (“blue”). Per-individual membership proportions for the blue cluster varied among Pygmy populations from 0.173±0.103 in the Central/Eastern Baka (CBK) to 0.527±0.096 in the Eastern Bongo (EBG). In contrast, among non-Pygmies, per-individual membership proportions for the red cluster were on average 0.197±0.093.

Our *STRUCTURE* results accord with previous studies [24-27,29,31,34,99,100] and can be interpreted as signals of asymmetric admixture between Central African Pygmies and non-Pygmies. In this view, appreciable membership in the blue “non-Pygmy” cluster among Pygmies reflects substantial and variable levels of non-Pygmy admixture, whereas low levels of membership in the red “Pygmy” cluster among non-Pygmies suggest low levels of Pygmy introgression into non-Pygmies.

### Correlation between non-Pygmy admixture and trait variation

Given the phenotypic and genetic differences observed between Pygmies and non-Pygmies in our sample set, we sought to establish whether genetic factors might underlie observed phenotypic patterns. If genetic differences between Pygmies and non-Pygmies contribute to their phenotypic differences, we might expect measurements for these traits to be correlated with per-individual levels of genetic admixture. We therefore separately investigated each trait’s correlation with per-individual membership proportions in the blue “non-Pygmy” *STRUCTURE* cluster at *K*=2 (**Figure 3B**; “non-Pygmy admixture” henceforth).

We observed a significant positive correlation between non-Pygmy admixture and adult standing height with the 76 males (Pearson *r*=0.585, *P*=2.90×10^−8^; **Figure 4A**) and the 57 females (*r*=0.485, *P*=1.30×10^−4^; **Figure 4B**) with standing height data available among the 406 unrelated individuals in our genotype dataset. The correlations remained significant when restricted to the 61 male and 31 female Pygmy individuals (*r*=0.311 with *P*=0.015 and *r*=0.441 with *P*=0.013, respectively). These findings are in agreement with previous studies on the relationship between levels of non-Pygmy admixture and adult standing height in Central African Pygmies [32-34] and provide further support for an appreciable genetic component to the determination of body size differences between Pygmies and non-Pygmies.

**Figure 4.**
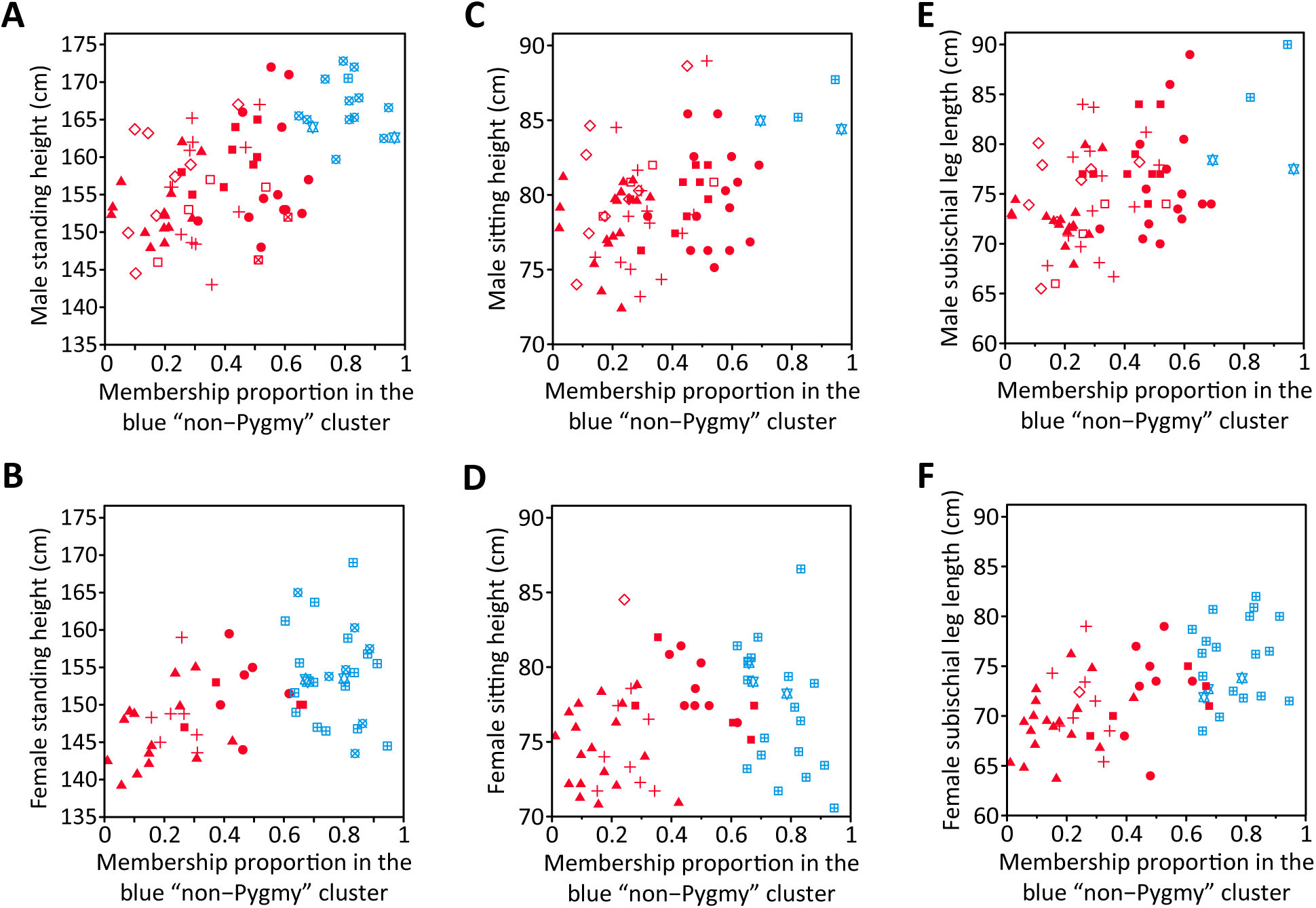
Relationship between non-Pygmy admixture and adult height-related traits. Scatterplots are shown of membership proportions in the blue “non-Pygmy” *STRUCTURE* cluster at *K*=2 (**Figure 3B**) and standing height for (**A**) 76 males (*r*=0.585, *P*=3×10^−8^) and (**B**) 57 females (*r*=0.485, *P*=3×10^−5^), sitting height for (**C**) 71 males (*r*=0.431, *P*=8.90×10^−5^) and (**D**) 57 females (*r*=0.226, *P*=0.040), and subischial leg length for (**E**) 71 males (*r*=0.475, *P*=1.41×10^−5^) and (**F**) 57 females (*r*=0.574, *P*=6.53×10^−7^). Only individuals included in the genetic dataset with a measurement for that trait were included in each comparison. Individuals are indicated by the symbols in **Figure 3A**.

The correlations of adult sitting height and subischial leg length with non-Pygmy admixture were similar in males (Pearson *r*=0.431 with *P*=8.90×10^−5^ and *r*=0.475, *P*=1.41×10^−5^, respectively; **Figures 4C** and **4E**). However, among females the correlation of non-Pygmy admixture with adult sitting height was markedly lower (*r*=0.226, *P*=0.040; **Figure 4C**) than with subischial leg length (*r*=0.574, *P*=6.53×10^−7^; **Figure 4F**). Considered together, these findings are consistent with common and distinct genetic factors contributing to sitting height and subischial leg length variation among Central Africans, while male– female differences in the strength of the correlations with non-Pygmy admixture might perhaps reflect the differential contribution of genetic variants in estrogen-dependent growth pathways to the determination of trunk and leg length [101,102].

Finally, while BMI was significantly positively correlated with non-Pygmy admixture in females (*r*=0.262, *P*=0.015; **Figure 5B**), BMI was only marginally so in males (*r*=0.166, *P*=0.060; **Figure 5A**). These correlations were markedly weaker than those obtained with the height-related measures (**Figure 4**), in accordance with smaller Pygmy/non-Pygmy differences in BMI (**Figure 2C**) than in adult standing height (**Figure 2A**) and compatible with non-genetic factors contributing substantially to body weight differences between Central African Pygmies and non-Pygmies.

**Figure 5.**
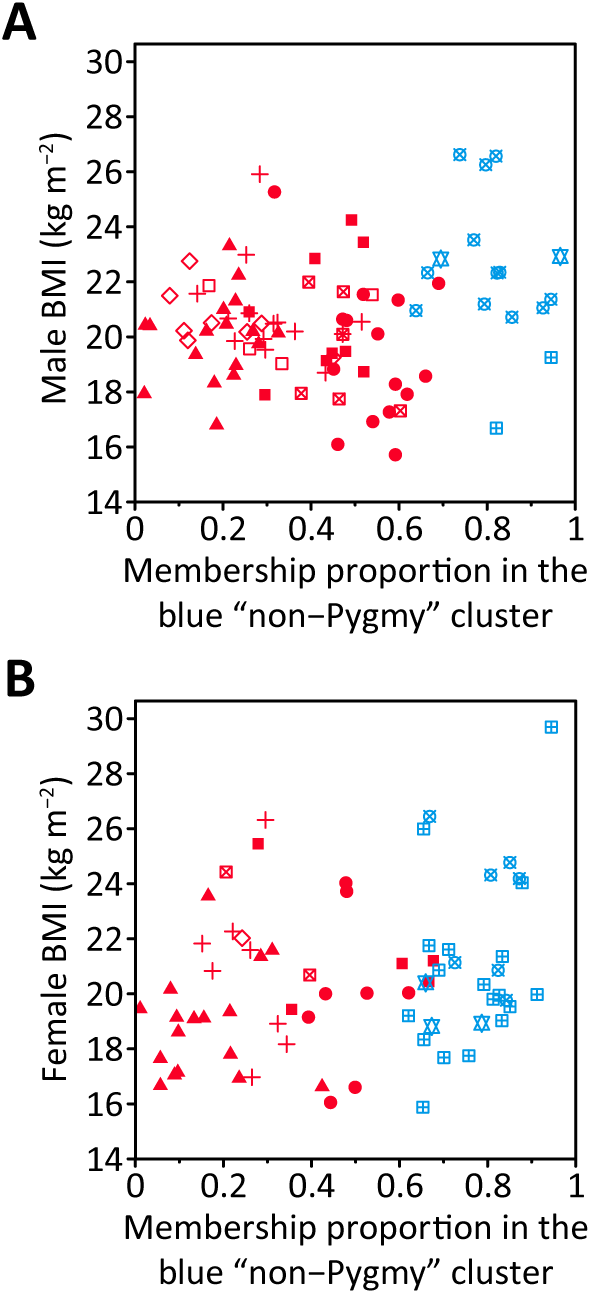
Relationship between non-Pygmy admixture and body mass index. Scatterplots are shown of membership proportions in the blue “non-Pygmy” *STRUCTURE* cluster at *K*=2 (**Figure 3B**) and BMI for (**A**) 89 males (*r*=0.166, *P*=0.060) and (**B**) 69 females (*r*=0.262, *P*=0.015) with both standing height and body weight measurements available in the genetic dataset were included in the comparison. Individuals are indicated by the symbols in **Figure 3A**.

These findings are consistent with a significant genetic component in the determination of height-related differences between Central African Pygmies and non-Pygmies. However, our results do not provide similar support for an appreciable genetic component to BMI variation patterns in Pygmies and non-Pygmies. Consequently, we next sought to identify genetic factors correlated with variation in height-related traits in our sample set of Pygmies and non-Pygmies, but not with variation in BMI.

### Association analyses of Pygmy/non-Pygmy categorization

We have found significant genetic and phenotypic differences between the Central African Pygmies and non-Pygmies in our sample set, with correlations between individual trait variation and inferred levels of non-Pygmy genetic admixture supporting an appreciable genetic component to the determination of their observed phenotypic differences. To investigate the underlying genetic component of their phenotypic differences, we first performed single-and multi-marker association tests to identify genomic regions harboring SNPs with significant allele frequency differences between the Pygmies and non-Pygmies in our genetic dataset of 406 unrelated individuals.

#### Single-marker association tests

To account for genetic structure (**Figure 3**) and cryptic genetic relatedness among individuals—which could inflate type-1 and type-2 errors [103-105]—in our single-marker association tests, we used *EMMAX*, which implements a linear mixed-effect regression model that corrects per-SNP association tests for structure and relatedness via a pairwise kinship matrix [105], and included village affiliation as a covariate (see **Materials and Methods**). We identified ten SNPs exhibiting a significant allele frequency differences between Pygmies and non-Pygmies after Bonferroni correction for multiple testing (α=5%), all lying within a single genomic region on chromosome 2 (**Figure 6C**). In addition, two further SNPs were marginally significant (α=10%; **Figure 6A** and **Table 2**), located on chromosomes 1 (Additional File 2: **Figure S2A**) and 7 (Additional File 2: **Figure S2B**). Our approach appeared to reasonably correct for structure and relatedness among individuals, as judged by the quantile-quantile plot (**Figure 6B**), while marginally inflated *P*-values were instead observed if village affiliation was not included as a covariate in the association tests (Additional File 2: **Figure S3**).

**Figure 6.**
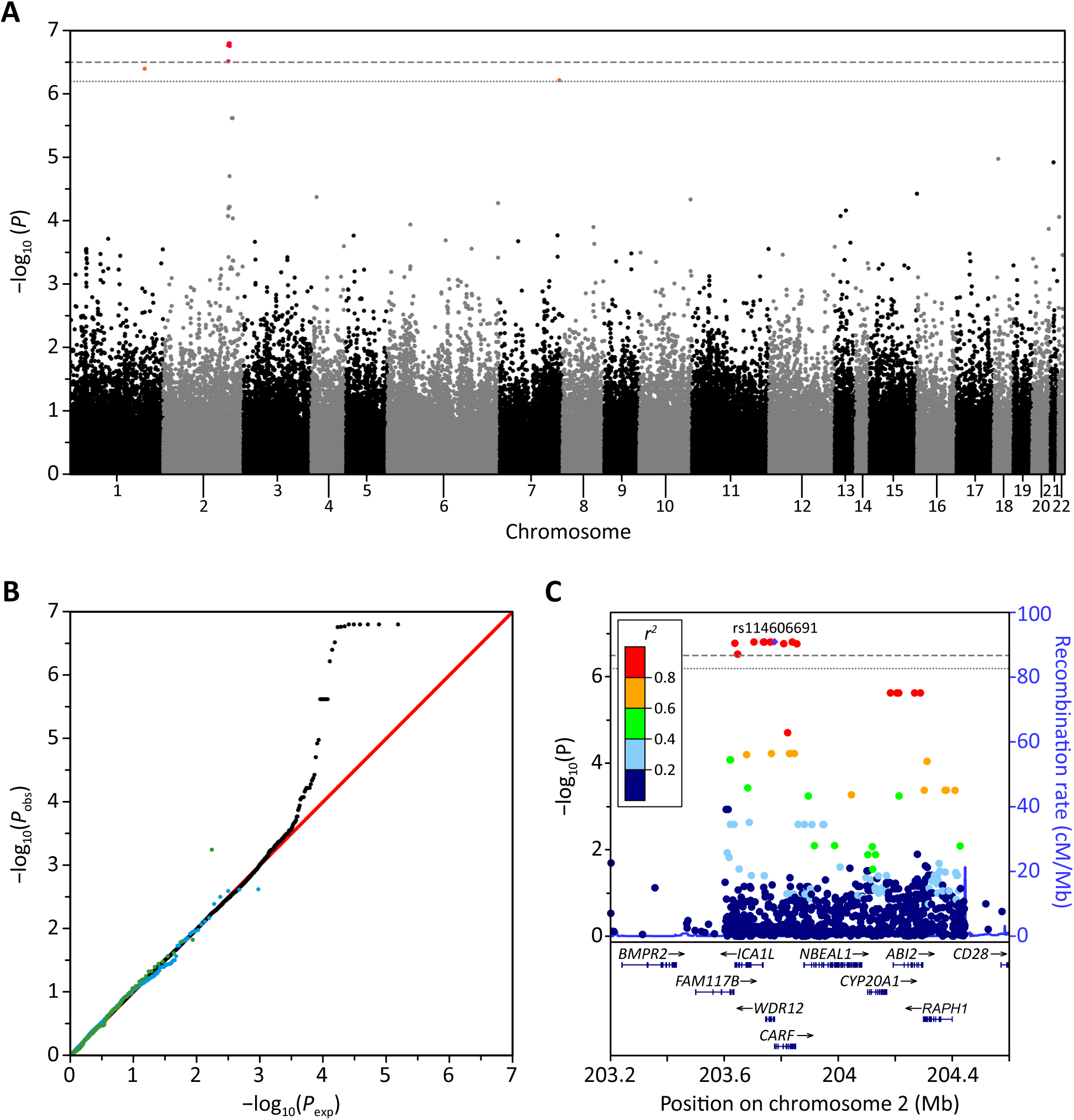
Association signals for Pygmy/non-Pygmy categorization. (**A**) Manhattan, and (**B**) quantile-quantile plots of the 153,798 autosomal SNPs in the genetic dataset. In **A**, the gray horizontal dashed line indicates the significance level for a Bonferroni correction at the 5% significance level while the dotted gray line indicates the significance level for a Bonferroni correction at the 10% significance level. In **B**, the identity line is shown in red, while the 126 SNPs identified by Wood *et al.* as significantly associated with standing height variation in Europeans [51] are plotted separately in green and the 949 SNPs listed in the Cardio-MetaboChip manifest [69] as associated with standing height variation are plotted separately in blue. (**C**) LocusZoom plot [147] of the genomic region surrounding the ten significant SNPs (**Table 2**). Top,-log_10_-transformed *P* values at individual SNPs colored by the strength of their correlation (*r^2^*) with the most significant SNP in the region (purple diamond), and HapMap Phase 2 recombination rates [148] depicted by the blue line. Bottom, gene locations in release hg19 of the UCSC database [145].

**Table 2.**
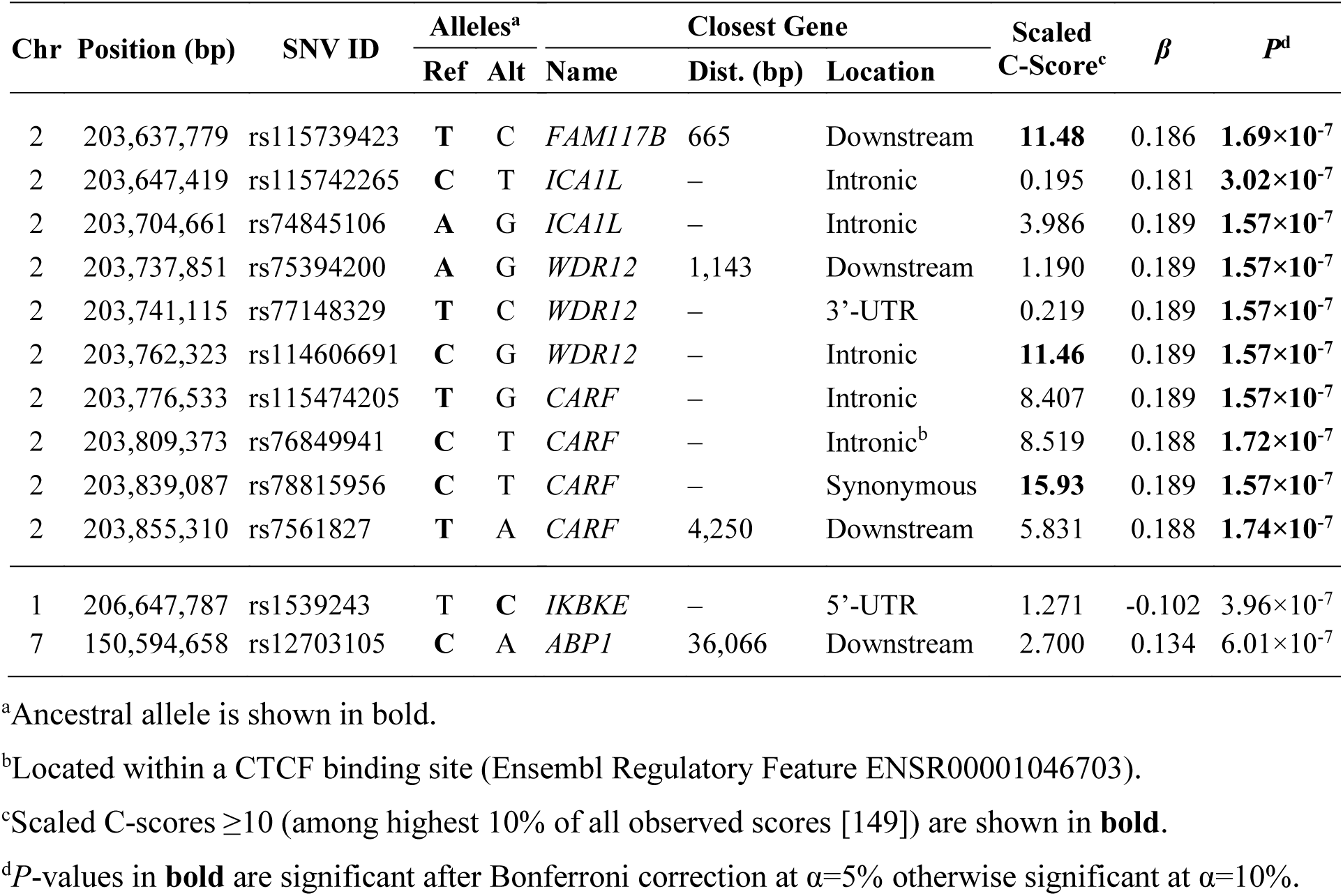
12 SNPs significantly associated with Pygmy/non-Pygmy categorization.

#### Multi-marker gene-based association tests

Because individual SNPs might fail to achieve the Bonferroni significance threshold owing to insufficient power in our modestly sized genetic dataset, we performed multi-marker gene-based association tests using *VEGAS* [106], a multivariate method that combines association signals across all SNPs located within a gene, correcting for non-independence between SNPs. We observed seven genes exhibiting a significant association with Pygmy/non-Pygmy categorization after Bonferroni correction (α=1%; **Table 3**); the genes identified did not differ appreciably if village affiliation was excluded as a covariate (Additional File 1: **Table S2**).

**Table 3.**
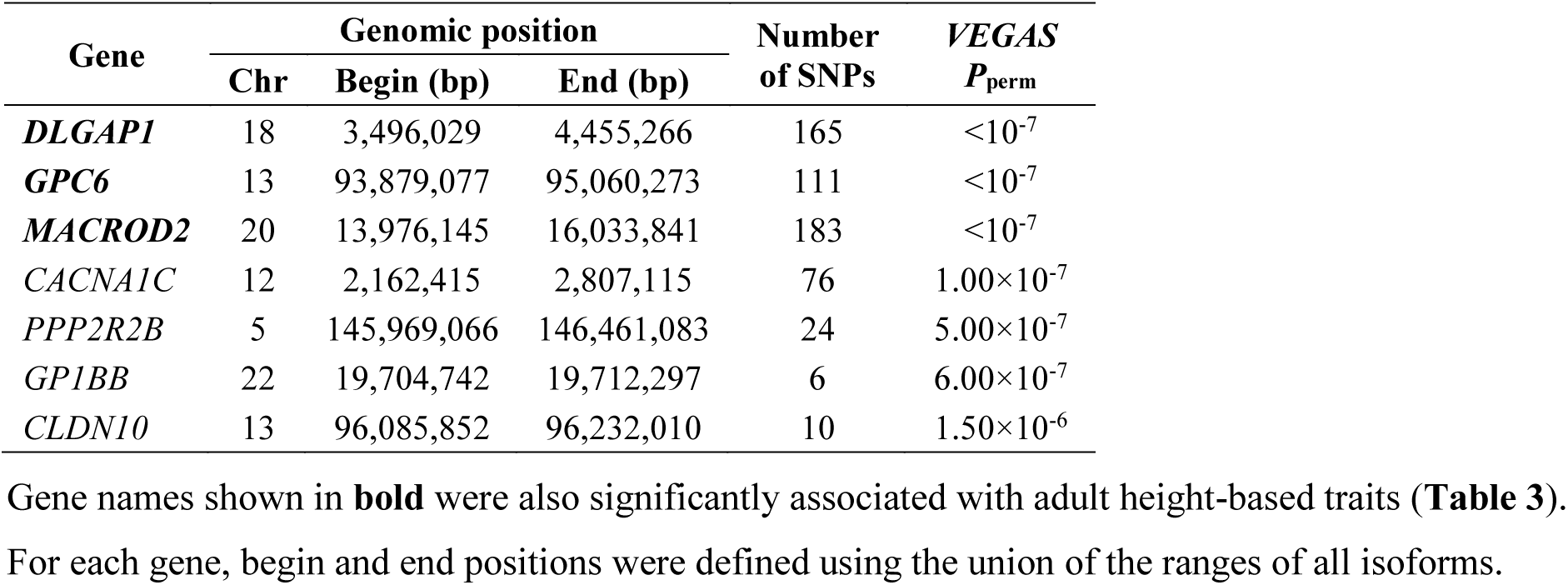
Seven genes significantly associated with Pygmy/non-Pygmy categorization.

#### Relevance to the Central African pygmy phenotype

While none of the genes in the genomic regions encompassing the ten significant SNPs (**Figure 6C**) and the two marginally significant SNPs (Additional File 2: **Figure S2**) had putative functions consistent with previously reported phenotypic differences between Central African Pygmies and non-Pygmies, one of the seven genes identified in the multi-marker analysis had a putative function in limb development. *GPC6* is a growth factor receptor that is important for correct growth plate formation [107] whose abrogation has been observed to cause long-bone growth retardation [108], suggesting that GPC6 is important for their longitudinal growth. This association is compatible with the observation that Pygmies have significantly shorter legs compared with non-Pygmies (**Figure 2B**) that are proportionally more reduced relative to trunk height (**Figure 2D**).

### Association analyses of height-related traits

Despite our modest sample size, we have identified genomic regions associated with an individual’s categorization as Pygmy or non-Pygmy that include one gene with a putative function compatible with their observed differences in leg length. Building upon this success, we next sought to identify genomic regions containing SNPs significantly correlated with variation in the height-related traits that differed significantly between the Pygmies and non-Pygmies in our sample set (**Figure 2A-D**).

#### Single-marker association tests

We performed per-SNP association tests to identify genomic regions harboring genes that contribute to variation in adult standing and sitting height and their ratio as well as subischial leg length among the 132-159 individuals with these data available in our dataset (Additional File 1: **Table S1**) using *EMMAX* and including sex and Pygmy/non-Pygmy categorization as covariates. Again, *EMMAX* appeared to provide a reasonable correction for structure and relatedness in these analyses (Additional File 2: **Figures S4E-S4H**, respectively) while similar analyses controlling only for sex (Additional File 2: **Figures S4A-S4D**) or for sex and village affiliation (Additional File 2: **Figures S4I-S4L**) instead led to slightly inflated or deflated *P*-values, respectively.

We did not find any SNPs significantly associated with adult standing (Additional File 2: **Figure S5**) or sitting (Additional File 2: **Figure S6**) height or subischial leg length (Additional File 2: **Figure S7**) variation after Bonferroni corrections for multiple testing, likely reflecting insufficient power with our modest sample sizes. However, we did identify a SNP marginally associated with sitting/standing height ratio after Bonferroni correction for multiple testing (α=10%; **Figure 7A**).

**Figure 7.**
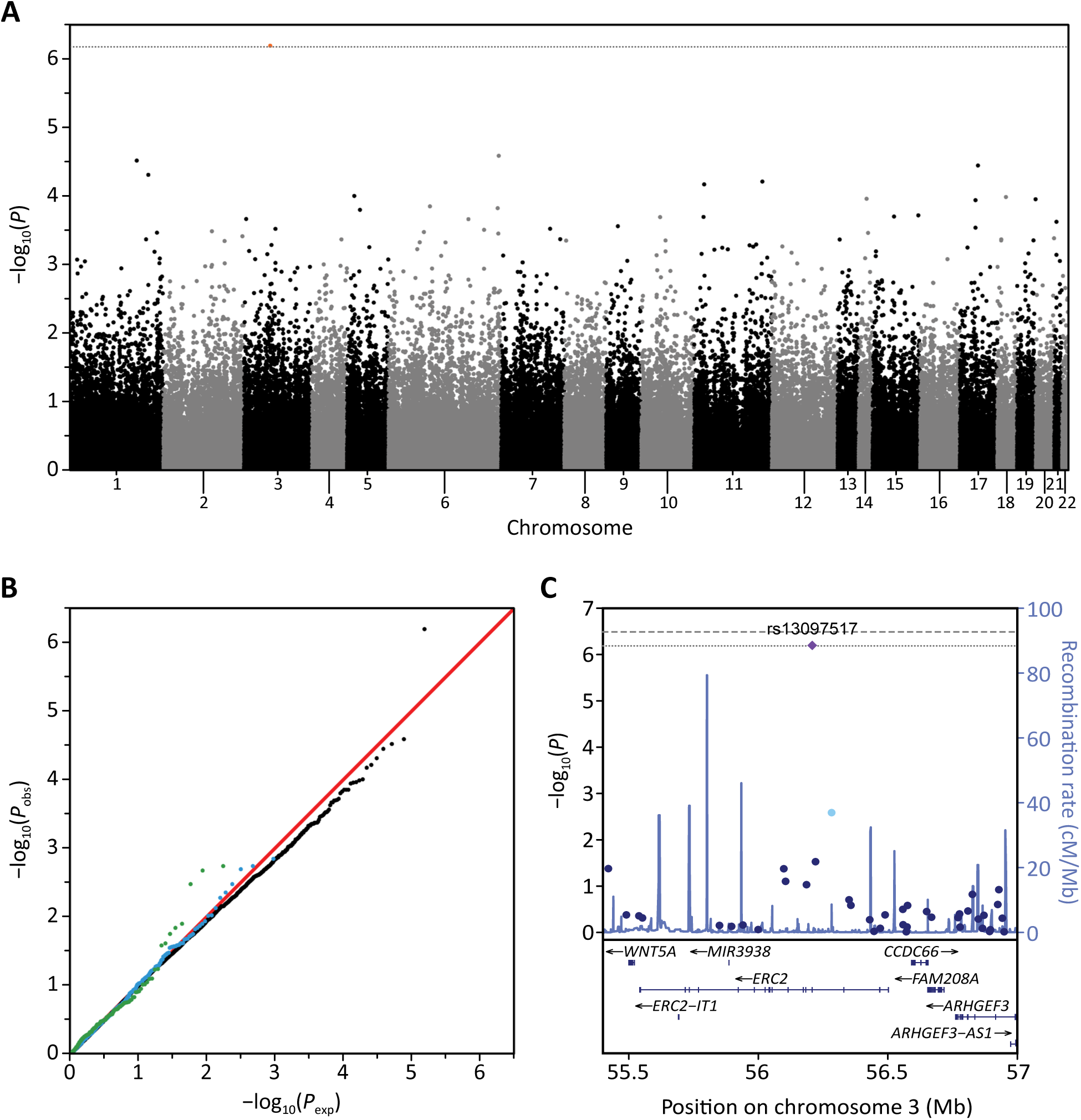
Association signals for adult sitting/standing height ratio. Manhattan (**A**) and quantile– quantile (**B**) plots of the 153,798 autosomal SNPs in the genetic dataset restricted to the 132 individuals with measurements for both standing and sitting height (Additional File 1: **Table S1**). (**C**) LocusZoom plot [147] of the genomic region surrounding the single marginally significant SNP (rs13097517, *P*=6.35×10^−7^). The figure follows the same format as **Figure 6**.

#### Multi-marker gene-based association tests

Gene-based association tests using *VEGAS* identified 19 genes associated with adult height-based traits after Bonferroni correction for multiple testing (α=1%; **Table 4**): nine with standing height, ten with sitting height, nine with sitting/standing ratio, and ten with subischial leg length. The genes identified for each trait did not differ markedly if only sex, or sex and village affiliation, were instead considered as covariates (Additional File 1: **Table S3**).

**Table 4.**
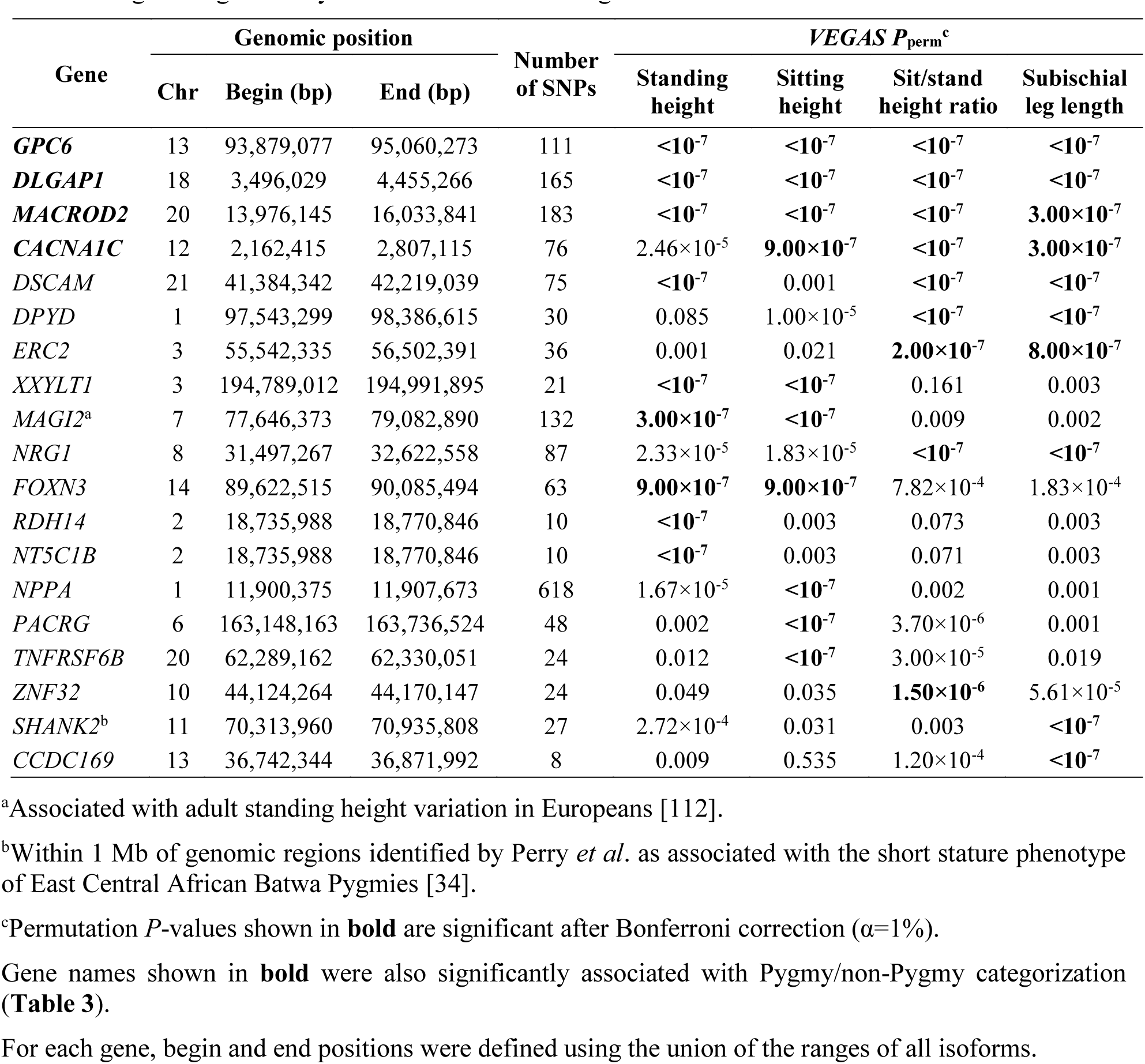
19 genes significantly associated with adult height.

#### Relevance to the Central African pygmy phenotype

The single SNP marginally associated with sitting/standing height ratio, rs13097517, is located in an intron of *ERC2* (**Figure 7C**), a member of the Rab3-interacting molecule (RIM)-binding protein family that functions as a regulator of neurotransmitter release [109], which lies ~600 kb upstream of *WNT5A*, a signaling ligand whose loss-of-function causes craniofacial and skeletal abnormalities including shortened limbs [110,111]. Thus, this association might reflect the influence of an untyped variant in strong LD with rs13097517 that modifies the expression of *WNT5A*, and potentially contributing to the differential skull morphology [59,62] and leg and forearm lengths [59,63] of Pygmies compared with non-Pygmies.

Three of the 19 genes were associated with all four height-based traits and were also identified in our *VEGAS* analysis for Pygmy/non-Pygmy categorization (**Table 3**), of which one (*GPC6*) has been implicated in the longitudinal growth of long bones [108]. In addition, *ERC2*, which is associated with both standing/sitting height ratio and subischial leg length (**Table 4**), contains SNP rs13097517, which was marginally associated with standing/sitting height ratio in our SNP-based association tests (**Figure 7C**). Importantly, one gene significantly associated with standing and sitting height variation in our Central Africans data set (*MAGI2*) has also been associated with standing height variation in non-Africans [112], while another gene significantly associated with subischial leg length (*SHANK2*) lies within a genomic region previously associated with the pygmy phenotype in Eastern Batwa Pygmies [34].

## Discussion

We have performed the first large-scale joint investigation of the variability and genetic architecture of the major components of adult standing height in Central African hunter-gatherer Pygmies and their agriculturalist non-Pygmy neighbors. Our findings accord with prior observations [59,63] that while Pygmy body size is generally proportionally reduced relative to non-Pygmies, their leg lengths are significantly shorter relative to their trunk length. Importantly, our results provide further support for an appreciable genetic component to the determination of body size differences between Pygmies and non-Pygmies, as implied by the correlations observed between the different measures and inferred levels of non-Pygmy admixture that replicate those reported previously for adult standing height [32-34].

Although our sample size is modest compared with traditional GWAS frequently conducted within populations of mainly European and Asian ancestry, our large-scale genetic association analyses using single-and multi-marker approaches identified three genomic regions as well as seven genes significantly associated with Pygmy/non-Pygmy categorization after a conservative Bonferroni correction for multiple testing. However, we were only able to identify a single genomic region associated with just one of the measured traits we considered (**Figure 7**). Given that for each trait a number of SNPs had notably lower *P*-values than the vast majority of those tested (Additional File 2: **Figures S5-S7**), our inability to identify additional genomic regions associated with variation in these traits likely reflects insufficient power with the much smaller sample sizes (132-159) than were available for our comparison of Pygmy/non-Pygmy categorization (406). Nevertheless, gene-based multi-marker test identified nine, ten, nine, and ten genes enriched for SNPs exhibiting an association with standing height, sitting height, sitting/standing height ratio, and subischial leg length, respectively. Importantly, quantile-quantile plots of the *P*-values created in each association analysis (Additional File 2: **Figures S3 and S4**) indicate that our approach appropriately corrected for structure and relatedness present among individuals in our dataset (**Figure 3**). Considered together with our identification of significantly associated genes and genomic regions, this would suggest that cross-population GWAS are a viable and powerful approach to identify the genetic basis of human phenotypes that naturally vary across populations.

A number of genes and genomic regions identified here potentially provide novel insight into the genetic basis of anatomical and physiological features of the Central African pygmy phenotype. Firstly, a number of genes associated with the height-related measures are reported to cause both skeletal and craniofacial abnormalities when perturbed: *MACROD2* [113,114], *GP1BB* [115,116], and *CACNA1C* [117,118]. It is therefore possible that craniofacial differences observed between Pygmies and non-Pygmies [59,62] may in part reflect the coevolution of these traits with Pygmy short stature; a hypothesis that remains to be formally tested in the absence of paleoanthropological data for Central Africa [3].

Secondly, the sole SNP marginally associated with sitting/standing height ratio (rs13097517) lies ~619 kb upstream of *WNT5A*, a signaling ligand whose abrogation causes craniofacial and skeletal abnormalities including shortened limbs [110,111]. Thus, in the context of the significantly higher sitting/standing height ratios we (**Figure 2D**) and others [59,63] have observed in Pygmies compared with non-Pygmies, this association would be compatible with a scenario in which Pygmy short stature is attributable in part to perturbation of *WNT5A* expression by one or more genetic variants in high LD with rs13097517. Intriguingly, WNT5A mediates the actions of growth factor receptor GPC6 [119], an important determinant of long bone growth [107,108] that was associated with Pygmy/non-Pygmy categorization (**Table 3**) and all four height-related traits (**Table 4**), suggesting that perturbation of the GPC6/WNT5A pathway may partly underlie Pygmy short stature.

Finally, a number of genes associated with Pygmy-non-Pygmy categorization have functions in immune rather than developmental processes. For example, *IKBKE* is an essential regulator of antiviral signaling pathways [120,121] and *CAPN2* is implicated in immune synapse formation in T-cell receptor signaling [122], while *FCGR1A* functions in monocyte phagocytosis [123] and has been implicated in dengue virus infection [124], a mosquito-borne disease present throughout Central Africa [125]. Our findings therefore add to the expanding body of evidence suggesting that differential adaptation in both immune-and auxologic-related processes have contributed to the evolution of the Central African pygmy phenotype [33,34,126].

Consistent with prior studies that investigated the contributions of >100 non-African standing-height-associated SNPs to standing height variation among 5 to 230 Central African Pygmies and non-Pygmies [33,34,126], none of the 949 SNPs listed in the Cardio-MetaboChip manifest [69] as associated with adult standing height were found to be individually associated with the height-related variation in our Central African sample set. Indeed, *P*-values for the 173 standing height SNPs reported by Wood *et al*. [51] and the 949 standing-height SNPs included on the Cardio-MetaboChip [69] based on the findings of Lango Allen *et al*. [45] did not depart from the uniform expectation to a greater extent than non-standing-height SNPs in our analyses of Pygmy/non-Pygmy categorization (**Figure 6B**) and subischial leg length (Additional File 2: **Figure S7B**). However, the Wood *et al*. SNPs did deviate to a greater extent in our standing height (Additional File 2: **Figure S5B**) and sitting/standing height ratio (**Figure 7B**) analyses while both the Wood *et al*. and Lango Allen *et al*. SNPs did deviated to a greater extent in our sitting height analysis (Additional File 2: **Figure S6B**). These observations indicate that previously reported non-African standing-height SNPs predominantly reflect perturbations of pathways contributing to trunk height but not to leg length. In contrast, the genes identified in our analysis of different height measures in Central Africans instead appear predominantly to reflect perturbation of pathways contributing to the determination of leg length. Our analyses therefore highlight the importance of conducting studies in Central African populations that can contribute novel information about the determinants of human standing height and body size proportions in general.

Despite our inability to detect SNP-wise associations with non-African standing height SNPs, gene-based association tests did identify one non-African standing height gene as being associated with standing and sitting height variation in Central Africans, *MAGI2* [112]. Thus, though the non-African standing height associated SNPs failed individually to reach significance in our association analyses, our identification of *MAGI2* using a multi-marker approach suggests that this is at least partly a consequence of insufficient power in our sample set. However, we cannot discount the possibility that for a subset of the non-African SNPs the absence of a significant association may be a result of discordance in LD patterns between Europeans and Central Africans disrupting the linkage between the genotyped SNP and the trait-determining variant [127].

## Conclusions

Despite the modest sample size in our study, we have identified genomic regions encompassing genes that are biologically meaningful in the context of both the traits themselves and phenotypic differences that exist between Central African Pygmies and non-Pygmies. Further, our findings provide additional support for an appreciable genetic component to the determination of phenotypic differences between Central African Pygmies and non-Pygmies. More generally, our results highlight the need for joint analyses considering different anthropometric and physiological measures in larger samples of Central African Pygmies and non-Pygmies to facilitate our understanding of the biological basis of the Central African pygmy phenotype. Future large-scale studies of worldwide short-stature populations have the potential to shed light on relative contributions of shared and distinct pathways in the development and maintenance of short stature. In particular, studying populations of different stature living in different ecologic environments will provide a greater understanding of adaptive processes underlying the appreciable variation in adult height observed across contemporary worldwide human populations.

## Material and Methods

### Samples

This study was conducted according to ethical principles of the Declaration of Helsinki. IRB approvals were obtained from the French Ministry of Higher Education and Research, University of Michigan, Stanford University, Washington State University and the University of Manitoba. Prior to sample collection, research authorizations were obtained from the Ministry of Public Health in Cameroon, the Ministry of Higher Education and Research in Gabon, the National Council for Science and Technology in Uganda, and the Ministry of Scientific Research in CAR, and informed consent was obtained from all research participants. In total, our study included 558 individuals from 20 Central African populations (**Figure 1**, **Table 1**). We conducted ethnographic interviews at each sampling site to categorize sampled populations *a priori* as “Pygmy” or “non-Pygmy” based on historical and cultural criteria that do not include adult height [4,6,7,23,128-131]. A community was categorized as “Pygmy” when it: (1) is recognized by outsiders as specialized in forest activities such as hunting-gathering and medical and magical knowledge of the rainforest; (2) shares complex socio-economic relationships with specific neighboring outsiders, such as exchanging forest products (e.g. game, wild honey) for iron tools (e.g. fishing hooks, iron blades, or spear heads); (3) is designated as “Pygmy” or its literal local translation or at least as “other than self” by neighboring populations; (4) distinguishes itself as a community with a different ethno-name from other neighboring communities, regardless of languages spoken; (5) has differing musical practices and instruments recognized as such by neighbors.

For each individual, DNA was extracted from either whole blood buffy coats with the DNeasy Blood & Tissue spin-column Kits (Qiagen, Valencia, CA), or saliva collected using Oragene kits (DNA Genotek Inc., Kanata, ON), following the manufacturer-recommended protocol. For a subset of the individuals from seven Pygmy—Baka (Center), Baka (East), Bezan (South), Bongo (Center), Bongo (East), Bongo (South), and Koya—and three non-Pygmy—Nzime, Tikar, and Bangando—populations, standing and sitting height were measured with a height gauge and weight with a standard weigh scale following standard anthropometric procedures [132]. Subischial leg length was calculated as standing minus sitting height, while BMI was calculated as body weight in kilograms divided by standing height in meters squared (kg m^−2^). Accurate age data was unavailable for these individuals, as most of the communities do not keep track of birth dates. Though we cannot rule out the confounding effect of osteoporotic age-related shrinking in our study, we expect it to be minimal as our sample set includes only adults and elderly individuals were not considered.

### Genotyping and quality control

DNA samples for 576 samples—558 individuals from 20 Central African Pygmy and non-Pygmy populations (**Table 1**) and 18 control samples—were genotyped at the University of Michigan Medical School DNA Sequencing Core (Ann Arbor, MI) using the Illumina MetaboChip that interrogates 196,725 genome-wide SNPs [69]. We focus on 196,091 SNPs whose genomic position had been independently verified (Peter Chines, unpublished data). Genotype calling was performed using the GenomeStudio Genotyping Module (v.1.0; Illumina Inc., San Diego, CA). Following quality control procedures conducted at the genotype-calling level (Stage 1, **Figure S8** [Additional File 2]), the preliminary dataset contained 192,903 autosomal SNPs that were polymorphic in a sample of 543 Pygmy and non-Pygmy individuals. Next, we conducted population-level quality control procedures (Stage 2, **Figure S8** [Additional File 2]), creating an initial dataset of 154,106 autosomal SNPs polymorphic in a sample of 543 Pygmy and non-Pygmy individuals. Hardy-Weinberg equilibrium was evaluated separately in each population using Yates-corrected chi-squared tests [133], and we used the same exclusion criteria as in Pemberton *et al*. [134].

Pygmy and non-Pygmy neighbors interact socially and economically on a daily basis [6,7,130,135]. Thus, individuals from neighboring communities might be present at the time of sampling. On rare occasions, some non-Pygmies have fled their communities (to avoid taxes or military recruitment for instance) and taken refuge among neighboring Pygmy populations. Therefore, it is possible that some sampled individuals were wrongly categorized according to our Pygmy/non-Pygmy categorization criteria. To search for categorization errors, we performed MDS analysis of individual pairwise allele-sharing distances (see below) in our initial dataset. We identified 16 individuals—11 initially categorized as Pygmies and five as non-Pygmies—who did not cluster genetically with other individuals sharing the same categorization (data not shown) and who might have been miscategorized during sample collection. To be conservative, we removed these 16 putative misclassified individuals from the preliminary dataset, and repeated the population-level quality control procedures (Stage 3, **Figure S8** [Additional File 2]) to create a dataset containing 154,029 autosomal SNPs polymorphic in a sample of 527 Pygmy and non-Pygmy individuals (“527RELAT” henceforth; **Table 1**); all individuals possessed genotypes at ≥94.9% of SNPs.

Relatedness among all pairs of the 527 individuals in the 527RELAT dataset was evaluated using identity-by-state allele sharing and the likelihood approach of *RELPAIR* (v.2.0.1) [136,137] following the methods of Pemberton *et al.* [134] restricted to sets of 9,999 SNPs. A total of 282 pairs of individuals were inferred by *RELPAIR* to be related at a level closer than first cousins: 230 intra-population and 52 inter-population pairs. All inter-population relative pairs involved geographically close populations (**Figure 1**); 47 involving the two Bezan populations (BZN and BZS), three involving two Bongo populations (CBG and EBG), and two with individuals from the nearby Fang (CFG) and Ngumba (NGB) populations. A dataset containing no first-or second-degree relatives was created by removing one individual from each of these 282 relative pairs. To minimize the number of individuals removed, we preferentially omitted individuals present in two or more relative pairs (either intra-or inter-population). In situations where either individual in a relative pair could be removed, we removed the individual with more missing data. After the exclusion of 121 related individuals—in addition to the 16 putative misclassified individuals—from the preliminary dataset of 543 individuals, many of whom were related to multiple individuals in the initial dataset, we repeated the population-level quality control procedures (Stage 4, **Figure S8** [Additional File 2]) to create an unrelated dataset with 153,798 autosomal SNPs polymorphic in a sample of 406 Pygmy and non-Pygmy individuals (“406UNRELAT” henceforth; **Table 1**); all individuals possessed genotypes at ≥94.9% of SNPs. Because of small sample sizes for the two Bezan populations (11 for BZN, 17 for BZS) in the 406UNRELAT dataset, we combined them into a single population (“Bezan” henceforth; BEZ, **Table 1**). Both the BZN and BZS samples belong to the same Bezan ethnic group located in Central Cameroon, which has a census size <400, divided between two nearby communities (~50 km apart) in frequent contact (P. Verdu and A. Froment, unpublished data).

### Population genetic analyses

#### Multidimensional scaling

We performed MDS based on ASD matrices constructed for all pairs of individuals in the 406UNRELAT dataset using *asd* (v1.0; https://github.com/szpiech/asd). This program considers in the calculation for a given pair only those SNPs for which neither individual was missing genotypes. We applied classic metric MDS on the ASD matrix using the *cmdscale* function in R (v2.11.0) [138]. To estimate the proportion of variation in ASD explained by the first two dimensions of the MDS plot, we calculated the Spearman correlation coefficient *ρ* between the Euclidean distances for all pairs of individuals and their corresponding ASD values (using *cor.test* in R here and for other correlations). To evaluate dispersion levels among Pygmies and non-Pygmies in the MDS plot, we compared the variance among their ASD values with a one-sided *F*-test using *var.test* in R.

##### STRUCTURE

To investigate genetic structure in our 406UNRELAT dataset, we performed model-based Bayesian clustering analyses implemented in *STRUCTURE* (v2.3) [97,98], which probabilistically assigns proportions of each individual’s genotypes to each of *K* genetic clusters, where *K* is set *a priori*, based on allele frequencies and irrespective of individual population categorizations. We used the *admixture* model with separate Dirichlet parameters in each cluster and correlated allele frequencies, and a burn-in period of 20,000 iterations followed by 10,000 iterations.

To minimize the number of linked loci owing to the gene-centric design of the Cardio-MetaboChip, in the *STRUCTURE* analyses we used 40,424 SNPs that have minimum spacing 18.75 kb. From these 40,424 SNPs, we created four non-overlapping panels of 10,106 SNPs, each with minimum marker spacing 75 kb. For SNP panel *n* (1 ≤ *n* ≤ 4), every fourth SNP along a vector of the considered 40,424 SNPs was selected, starting at position *n* – 1. In this vector, SNPs were numbered starting at 0 and ordered from chromosome 1 to 22 and by increasing distance along each chromosome (taking genomic positions from NCBI database build 37). For each panel, we computed 10 independent *STRUCTURE* runs for values of *K* between 2 and 4, producing 40 independent replicates for each *K.* We identified common modes among the 40 replicates using *CLUMPP* [139] with the *Greedy* algorithm and 1,000 random permutations. For each *K*, all pairs of runs with a symmetric similarity coefficient >0.9 were considered to belong to the same mode. For each mode, we computed individual membership proportions averaged across runs from that mode, visualizing the most frequent mode at each *K* using *DISTRUCT* [140].

To investigate the relationship between non-Pygmy admixture and adult height in Central Africans, we calculated the Pearson product-moment correlation coefficient *r* between individual membership proportions in the blue cluster at *K=2* and measured adult height where available.

### Association analyses

#### Single-marker tests

Per-SNP association tests were performed using *EMMAX* [105], which implements a mixed-effect regression model [141] to account for genetic structure in a sample set, incorporating variance components of random polygenic effects [142,143] and genetic relatedness among individuals using a pairwise Balding-Nichols kinship relatedness matrix [144] (equation 7 in [105]). We constructed the kinship matrix for all pairs of individuals in the 406UNRELAT dataset using *EMMAX*, including in the calculation only the 94,050 and 93,821 SNPs, respectively, inferred to be in linkage equilibrium (*r*^2^<0.5). Association analyses for Pygmy/non-Pygmy categorization were performed by labeling Pygmies as “cases” and non-Pygmies as “controls.” Association analyses were performed separately for each measured trait considering only those individuals with data available. To improve the normality of the body weight and BMI distributions and alleviate the impact of outliers, body weight and BMI values were rank-based inverse-normal transformed separately for each gender, and association tests performed using these transformed values. To control for population structure and sexual dimorphism in our sample set, in our analysis of Pygmy and non-Pygmy categorization we considered the residuals of the regression of Pygmy/non-Pygmy categorization on village affiliation, while for quantitative-traits we considered the residuals of a multiple regression for that trait on sex and Pygmy/non-Pygmy categories.

#### Multi-marker gene-based tests

As individual SNPs may fail to achieve the significance threshold due to insufficient power in our modest sample sizes, we performed multi-marker gene-based tests of association using *VEGAS* [106]. Separately for each gene, *VEGAS* computes a *χ*^2^ test statistic from observed *EMMAX P-values* for all SNPs that lie within its boundaries (defined as ±50 kb from the ends of the transcribed region) and evaluates significance via simulations from a multivariate normal distribution with mean 0 and a covariance matrix of pairwise LD between SNPs. The transcribed region of each gene was defined using release hg19 of the UCSC gene database [145]; for genes with multiple isoforms, begin and end positions were defined as the outermost positions in the union of the transcribed regions of all isoforms. Pairwise LD estimates in our datasets considered in the calculation only unrelated individuals.

## List of Abbreviations Used

AR: androgen receptor
ASD: allele-sharing dissimilarity
BMI: body mass index
DRC: Democratic Republic of Congo
GWAS: genome-wide association study
HGH: human growth hormone
IGF1: insulin-like growth factor 1
MDS: multidimensional scaling
SD: standard deviation
SNP: single nucleotide polymorphism.

## Acknowledgements

The authors thank the volunteers from Cameroon, Central African Republic, Gabon, and Uganda who participated in this study. The authors thank Peter Chines for providing verified Cardio-MetaboChip SNP positions, Sen Li for providing ancestral and derived allelic states for the MetaboChip SNPs, Hyun Min Kang for assistance with *EMMAX*, and Frédéric Austerlitz, Erkan Buzbas, Zachary Szpiech, and Lawrence Uricchio for useful comments and discussions.

## Declarations

**Ethics approval and consent to participate**

This study was approved by institutional ethics review committees at the French Ministry of Higher Education and Research, University of Michigan, Stanford University, Washington State University and the University of Manitoba. Research authorizations were obtained from the Ministry of Public Health in Cameroon, the Ministry of Higher Education and Research in Gabon, the National Council for Science and Technology in Uganda, and the Ministry of Scientific Research in CAR.

## Consent for publication

All authors read and approved the final manuscript.

## Availability of data and material

The Illumina Cardio-MetaboChip SNP genotype dataset for the 406 unrelated individuals analyzed in this study is available from the European Genome-Phenome archive (**<u><accession no. pending></u>**), and will be accessible for population and evolutionary history studies in accord with the informed consent documents used for this study.

## Authors’ contributions

EH, NAR, PV, TJP, NSB, and CJW conceived and designed the study. AF, BSH, PV, and SLB collected the samples. NSB prepared DNA samples for genotyping. PV performed the genotype calling and quality control with assistance from TJP and NSB. TJP performed data preparation and population-level quality control. PV performed population-genetic analyses with the assistance of TJP. TJP and PV designed and performed the association analyses. TJP, PV and NAR wrote the manuscript with the assistance of all other authors. All authors read and approved the final manuscript.

## Description of Additional Data Files

**Additional File 1:** Tables S1-S3

**Additional File 2:** Figures S1-S8

